# Plasma membrane nanoscale dynamics of *Arabidopsis* leucine-rich repeat receptor kinase complexes

**DOI:** 10.64898/2026.03.05.709869

**Authors:** Michelle von Arx, Marie-Dominique Jolivet, David Biermann, Vaishali Gabani, Steven S. Andrews, Cyril Zipfel, Julien Gronnier

## Abstract

Plasma membrane-localized receptors operate as dynamic signaling complexes and integrative networks^1–3^, yet the spatial and temporal regulation of these interactions remain largely unknown. Here, by analyzing the components of a minimal *Arabidopsis* leucine-rich repeat receptor kinase network, we describe the differential diffusion and organization of receptor complex components and unveil the nanoscale spatial and temporal logic underlying the formation of receptor kinase complexes. The ligand-binding receptors FLS2 and BRI1, and the accessory receptor BIR3, are organized in plasma membrane nanodomains, within which the co-receptor BAK1 diffuses and is spatially arrested upon ligand perception. BAK1’s spatial arrest relies on extracellular domain (ECD)-ECD interactions but does not require receptor complex activation. Mathematical modelling, single molecule imaging and bio-assays infer that accessory receptors maintain a dynamic pool of co-receptors in the vicinity of ligand-binding receptors to promote ligand-induced complex formation and signaling. We propose that ligand-induced receptor kinase complex formation is a deterministic process defined by the relative nanoscale spatial positioning of individual signaling and regulatory components.

## Main

Plants and metazoan cell-surface receptors sense and relay a myriad of extracellular signals to coordinate development and adjust to ever-changing environmental conditions^4,5^. In plants, the repertoire of cell-surface receptor kinases has considerably expanded with over 400 members encoded in the genome of *Arabidopsis thaliana* (hereafter Arabidopsis)^6^. Among them, leucine-rich repeat receptor kinases (LRR-RKs) constitute a prominent class that perceive self, non-self and modified-self molecules to regulate development, immunity and reproduction^7–9^. For instance, the ligand-binding LRR-RK BRASSINOSTEROID-INSENSITIVE 1 (BRI1) functions as the main receptor for plant steroid hormones brassinosteroids^7^ and forms a complex with the co-receptor BRI1-ASSOCIATED KINASE (BAK1, also called SOMATIC EMBRYOGENESIS RECEPTOR KINASE 3, SERK3) to regulate growth^7,10^. BAK1 also serves as a co-receptor for other LRR-RKs, such as FLAGELING SENSING 2 (FLS2) that sense the flg22 epitope derived from bacterial flagellin to initiate immune signaling^9,11^.

LRR-RKs have been proposed to function as a coordinated interaction network to process extracellular signals^2^. In this framework, accessory receptor kinases regulate ligand-induced association between ligand-binding receptors and their co-receptors. For instance, the LRR-RK BAK1-INTERACTING RECEPTOR-LIKE KINASE 3 (BIR3) physically associates with BAK1^12,13^ and has been proposed to inhibit ligand-induced FLS2-BAK1 and BRI1-BAK1 complex formation^13–15^. Several studies have highlighted the spatial organization of LRR-RKs into plasma membrane nanodomains^16–22^. For instance, the spatial separation of FLS2 and BRI1 has been proposed to support signaling specificity^16^ and alterations of the nanoscale organization of FLS2 and BAK1 correlate with defects in ligand-induced complex formation and signaling^17,23^. Yet, how the dynamic associations of cell surface receptor are coordinated in space and time within the plasma membrane remains unclear. Here, we studied a minimal LRR-RK network composed of the ligand-binding receptors FLS2 and BRI1, their common co-receptor BAK1 and the accessory receptor BIR3.

We analyzed the plasma membrane nanoscale organization of FLS2-GFP^24^, BRI1-GFP^25^ and BAK1-GFP^26^ by combining variable-angle total internal reflection microscopy (VA-TIRFM)^27,28^ with enhanced super-resolution radial fluctuations (eSRRF)^29^ in hypocotyl epidermal cells of light grown Arabidopsis seedlings. Consistent with previous observations made in Arabidopsis cotyledons epidermal cells^16–18,30,31^, we observed that FLS2-GFP and BRI1-GFP are organized into well-defined nanodomains (Fig. 1a-b). In contrast, BAK1-GFP showed a more dispersed distribution within the PM (Fig. 1a-b). To complement these observations, we used photochromic reversion, a variant of single-particle tracking photoactivated localization microscopy (spt-PALM) that enables long-term single molecule imaging^32^. We analyzed BRI1, FLS2 and BAK1 tagged with the photoconvertible fluorescence protein mEOS3.2^33^ expressed under the control of their native promoter. We observed that FLS2-mEOS3.2 and BRI1-mEOS3.2 are overall static while BAK1 is laterally diffusive and presents a higher diffusion coefficient (Fig. 1c-d; Extended Data Fig. 1a-b). We observed that when transiently expressed in *Nicotiana benthamiana,* or expressed under the control of the *ubiquitin 10* promoter^34^ in Arabidopsis stable transgenic lines, FLS2-mEOS3.2 remained static and BAK1 remained diffusive (Fig. 1c-d; Extended Data Fig. 1a-d, 2a, 3c). We next analyzed FLS2, BRI1 and BAK1 organization using nanoscale spatiotemporal indexing clustering (NASTIC)^35^. We observed that FLS2 and BRI1 are more clustered than BAK1 (Fig. 1e-f). Together these observations highlight the differential organization and diffusion of plant LRR-RK complex components, features that contrast with the behavior of animal cell-surface receptors^36–39^. We next investigated the effect of ligand perception on receptor dynamics. We imaged FLS2 and BRI1 within 10 min post ligand-treatment, a time-frame which corresponds to initiation of ligand-induced complex formation and signaling^11,40,41^. In good agreement with the analysis of FLS2-GFP diffusion in *N. benthamiana*^42^, we observed that flg22 does not lead to changes in FLS2-mEOS3.2 diffusion in *N. benthamiana* and in Arabidopsis (Fig. 2c-d; Extended Data Fig. 2a-b). Further, flg22 treatment did change FLS2 clustering (Extended Data Fig. 2c-d). Similarly, we observed that BRI1-mEOS3.2 nanoscale dynamics and clustering is unaffected by epi-brassinolide (eBL) treatment (Fig. 2a-b; Extended Data Fig. 2e-f). To corroborate these observations, we used computational analysis of spatial arrests (CASTA)^32^ to analyze changes in FLS2 and BRI1 dynamics in long-term single molecule imaging. We observed that flg22 and eBL do not lead to changes in FLS2 and BRI1 spatial arrest (Extended Data Fig. 2g-j). We conclude that ligand-induced complex formation and early ligand-induced signaling is not associated with changes in ligand-binding receptor nanoscale dynamics or organization.

**Figure 1.**
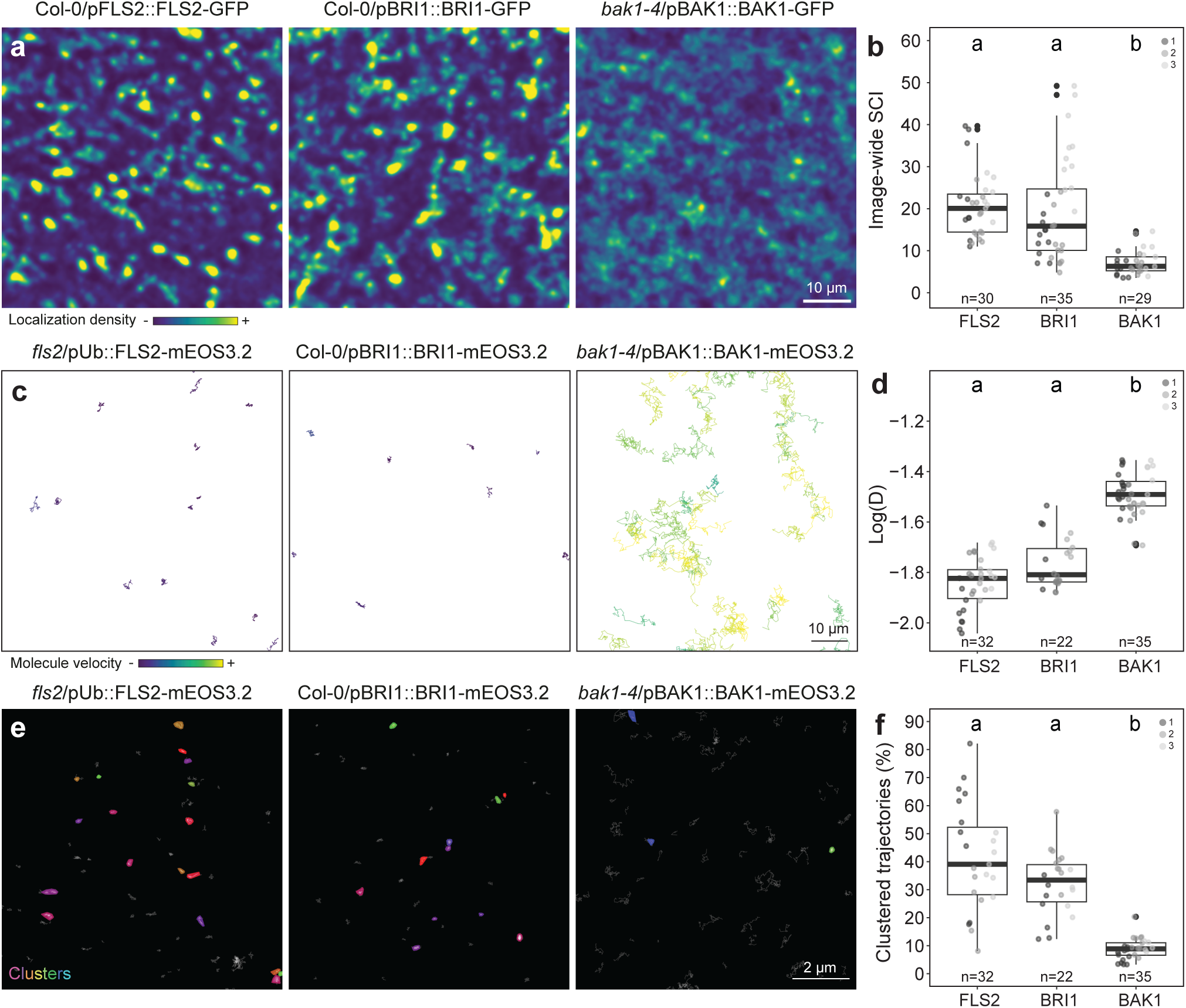
Differential organization and dynamics of LRR-RK complex components. **a**, Representative eSRRF images of FLS2-GFP, BRI1-GFP and BAK1-GFP plasma membrane organization in 5-day-old Arabidopsis hypocotyl. **b**, Quantification of the image wide spatial clustering index (iSCI). Each data point represents the average value per cell (n), colors indicate independent experiments. **c**, Representative images of FLS2-mEOS3.2, BRI1-mEOS3.2 and BAK1-mEOS3.2 long-term single molecule trajectories in 5-day-old Arabidopsis hypocotyl. Trajectories are colored based on velocity. **d**, Analysis of instantaneous diffusion coefficient (D). Each data point represents the average log(D) value in μm^2^/s obtained per cell (n), colors indicate independent experiments. **e**, Representative images of FLS2-mEOS3.2, BRI1-mEOS3.2 and BAK1-mEOS3.2 trajectories nanoclustering in 5-day-old Arabidopsis hypocotyl. **f**, Quantification of percentage of clustered trajectories. Each data point represents the percentage of clustered trajectories per cell (n), colors indicate independent experiments. Conditions sharing a letter are not significantly different in pairwise Wilcoxon with Bonferroni correction (p>0.05).

**Figure 2.**
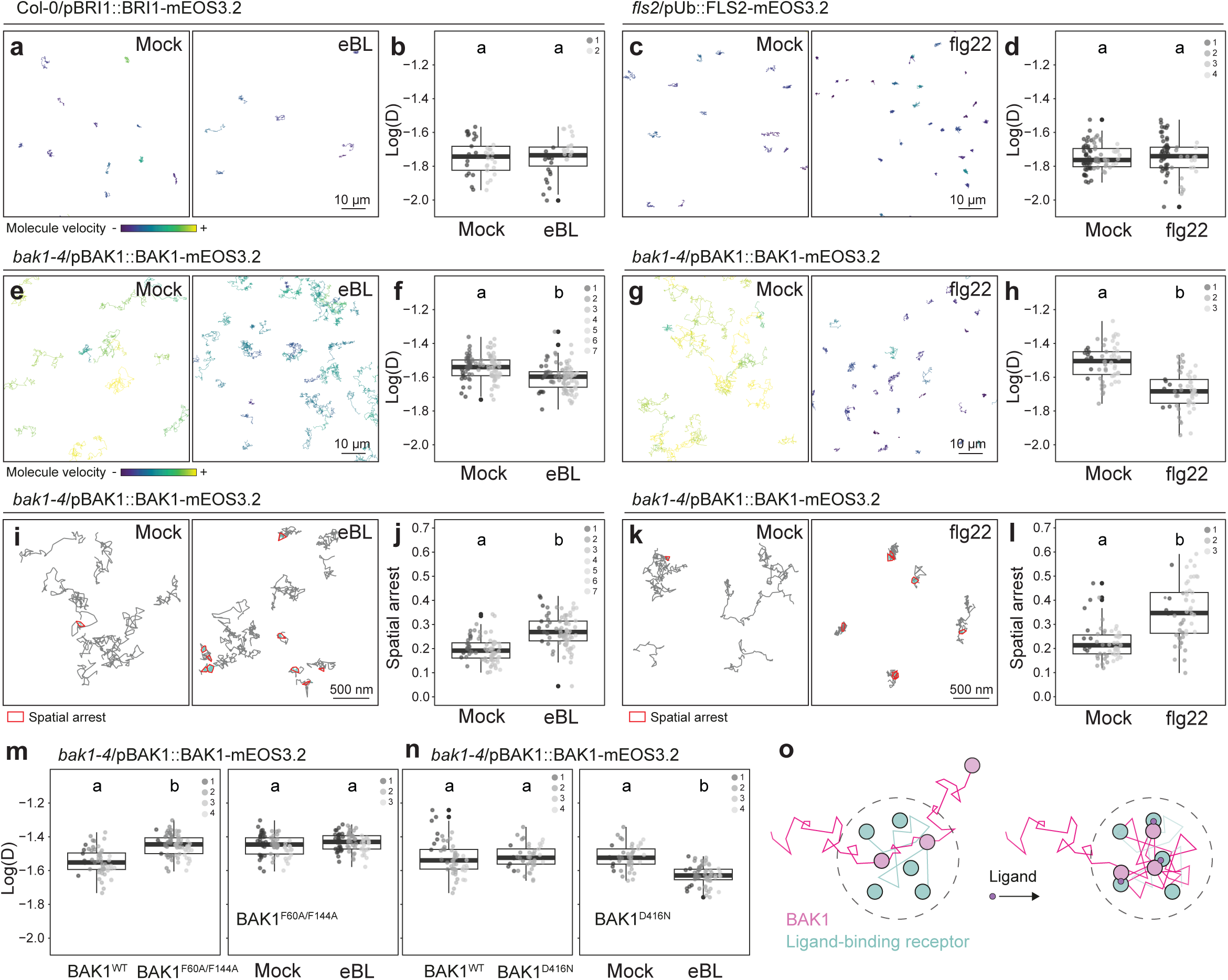
BAK1 is spatially arrested upon ligand perception. **a**, Representative BRI1-mEOS3.2 trajectories, long-term single molecule imaging in presence of 1 µM eBL or corresponding mock control (EtOH) in Arabidopsis hypocotyl. **b**, Quantification of instantaneous diffusion coefficient (D) within 10 min of treatment. Each data point represents the average log(D) value obtained per cell (n), colors indicate independent experiments. **c**, Representative FLS2-mEOS3.2 trajectories, long-term single molecule imaging in presence of 1 μM flg22 or corresponding mock control (H_2_O) in Arabidopsis hypocotyl. **d**, Quantification of instantaneous diffusion coefficient (D) within 10 min of treatment. Each data point represents the average log(D) value obtained per cell (n), colors indicate independent experiments. **e**, **g**, **i**, and **k**, Representative BAK1-mEOS3.2 trajectories, long-term single molecule imaging in presence of 1 µM eBL (**e** and **i**) or 1 μM flg22 (**g** and **k**) or the corresponding mock controls (EtOH or H_2_O) in *bak1-4* (**e** and **i**) and *bak1-4*/pFLS2::FLS2-GFP (**g** and **k**) hypocotyls. **f**, **h**, **m** and **n**, Quantification of instantaneous diffusion coefficient (D) within 10 min of treatment. Each data point represents the average log(D) value obtained per cell (n), colors indicate independent experiments. **j** and **l**, Quantification of BAK1-mEOS3.2 spatial arrests within 10 min of treatment. Each data point represents the average number of spatial arrests per trajectory obtained per cell (n), colors indicate independent experiments. Conditions sharing a letter are not significantly different in pairwise Wilcoxon test with Bonferroni correction (p>0.05). **o**, Working model: BAK1 navigates within and is stabilized to pre-formed ligand-binding RK nanodomains upon ligand perception.

We next examined the behavior of BAK1. We observed that BAK1-mEOS3.2 diffusion coefficient is reduced upon eBL treatment (Fig. 2e-f). Further, we observed an increase in BAK1-mEOS3.2 clustering (Extended Data Fig. 3a-b) and in the frequency of its spatial arrest (Fig. 2i-j) events upon eBL treatment. Similarly, we observed that flg22 treatment reduces BAK1-mEOS3.2 diffusion in *N. benthamiana* (Extended Data Fig. 3c-d). However, in Arabidopsis hypocotyl cells, we did not observe changes in BAK1 mobility behavior (Extended Data Fig. 3e-f). We presumed that the low FLS2 expression in Arabidopsis hypocotyl cells (Extended Data Fig. 1a) combined with the inherent stochasticity of BAK1-mEOS3.2 photoconversion impedes our ability to detect changes in BAK1 mobility behavior. To test this hypothesis, we crossed *bak1-4*/pBAK1::BAK1-mEOS3.2 with Col-0/pFLS2::FLS2-GFP to increase FLS2 accumulation. We observed that introducing FLS2-GFP in *bak1-4*/pBAK1::BAK1-mEOS3.2 leads to detectable flg22-induced decrease in BAK1-mEOS3.2 diffusion (Fig. 2g-h) and increase in BAK1 clustering (Extended Data Fig. 3g-h). Further, we observed that similar to eBL, flg22 treatments lead to an increase in BAK1-mEOS3.2 spatial arrests (Fig. 2i-l). Together these observations show that, while ligand binding RKs are static and organized in clusters independently of exogenous ligand treatment, BAK1 clustering and spatial arrestment increases upon ligand perception. Using dual-color VA-TIRFM we observed that FLS2-GFP and BAK1-mCherry, as well as BRI1-GFP and BAK1-mCherry partially co-localize in control conditions (Extended Data Fig. 4a-b and d-e). These observations are consistent with previous cell surface-selective FLIM analyses of BRI1-GFP and BAK1-mCherry^20^, and suggest that BAK1 navigates in the vicinity of ligand-binding receptor nanodomains. Further, we observed that loss of the receptor kinase FERONIA (allele *fer-*4), which induces an increase in BRI1-BAK1 association^43^, led to an increased co-localization between BRI1-GFP and BAK1-mCherry (Extended Data Fig. 4f). Similarly, we observed that flg22 treatment leads to an increased co-localization between FLS2-GFP and BAK1-mCherry (Extended Data Fig. 4c). In contrast to BRI1-BAK1, flg22-induced FLS2-BAK1 complex formation was reduced in *fer-4*^44^. In agreement, we observed flg22 treatment does not lead to an increase in co-localization between FLS2-GFP and BAK1-mCherry in *fer-4*. Together with single molecule tracking analyses, these observations suggest that BAK1 navigates within and is spatially arrested to pre-formed ligand-binding receptor nanodomains upon ligand perception.

We next investigated the molecular events underlying ligand-induced BAK1 spatial arrestment. We mutated two residues in the BAK1 ECD, Phenylalanine (F) 60 and F144, which are commonly engaged in interaction with ligand-bound ligand-binding receptors, including BRI1, and are required for ligand-induced complex formation^13,45–48^. We observed that the nanoscale dynamics and organization of BAK1^F60A/F144A^-mEOS3.2 is insensitive to eBL treatment (Fig. 2m; Extended Data Fig. 5a-c), suggesting that ligand-induced BAK1 spatial arrestment relates to ligand-induced complex formation. In agreement, using a BAK1 kinase-dead version of BAK1 (BAK1^D416N^)^49,50^, we observed that BAK1 kinase activity is not required for eBL-induced BAK1 spatial arrestment and clustering (Fig. 2n; Extended Data Fig. 5d-f). We conclude that ligand-induced BAK1 spatial arrestment corresponds to ligand-induced complex formation and does not rely on receptor complex activation and downstream signaling events.

We observed that a fraction of BAK1 is spatially arrested in absence of exogenous ligand treatment (Fig. 2) and that mutations that abolish BAK1 association with a variety of ligand-binding receptors (Fig. 2m) are not sufficient to abolish BAK1 clustering and spatial arrestment (Extended Data Fig. 5g-i). We thus hypothesized that additional regulators control BAK1 spatial arrestment. The accessory receptor kinase BIR3 physically associates with BAK1^13,14^ and is in proximity to FLS2 and BRI1^14^. Using VA-TIRFM-eSRRF we observed that, similar to FLS2 and BRI1 (Fig. 1), BIR3-GFP is organized into plasma membrane nanodomains (Extended data Fig. 6a). Corroborating these observations, we observed that BIR3-mEOS3.2 is relatively static and forms nanoclusters (Fig. 3a-d; Extended data Fig. 6b-c). Since BIR3 showed a slow and confined lateral diffusion, we hypothesized that BAK1 transiently physically associates with BIR3 and conditions BAK1 spatial arrestment. We therefore mutated BAK1 Aspartic acid (D) 122, which has been shown to reduce BAK1-BIR3 ECD-ECD interactions *in vitro*^13^. We observed that substituting D122 with Alanine (A) led to an increase in BAK1 diffusion (Fig. 3e-f), as well as a reduction in BAK1 spatial arrestment (Fig. 3g-h) and clustering (Fig. 3i-j), indicating that BIR3 regulates BAK1 nanoscale dynamics. Corroborating these observations, we observed that loss of *BIR3* leads to an increase in BAK1-mEOS3.2 diffusion and a decrease in BAK1 clustering (Extended Data Fig. 7a-e). Loss of *BIR3* was not sufficient to significantly affect BAK1 spatial arrest (Extended Data Fig 7f-g), suggesting the implication of additional factors; potentially the BIR3 paralogues BIR1 and/or BIR2^51,52^. We conclude that the association of BAK1 to BIR3, and potentially other BIRs, contribute to BAK1 clustering and spatial arrestment presumably in the vicinity of the ligand-binding receptor kinases FLS2 and BRI1 (Fig. 3k).

**Figure 3.**
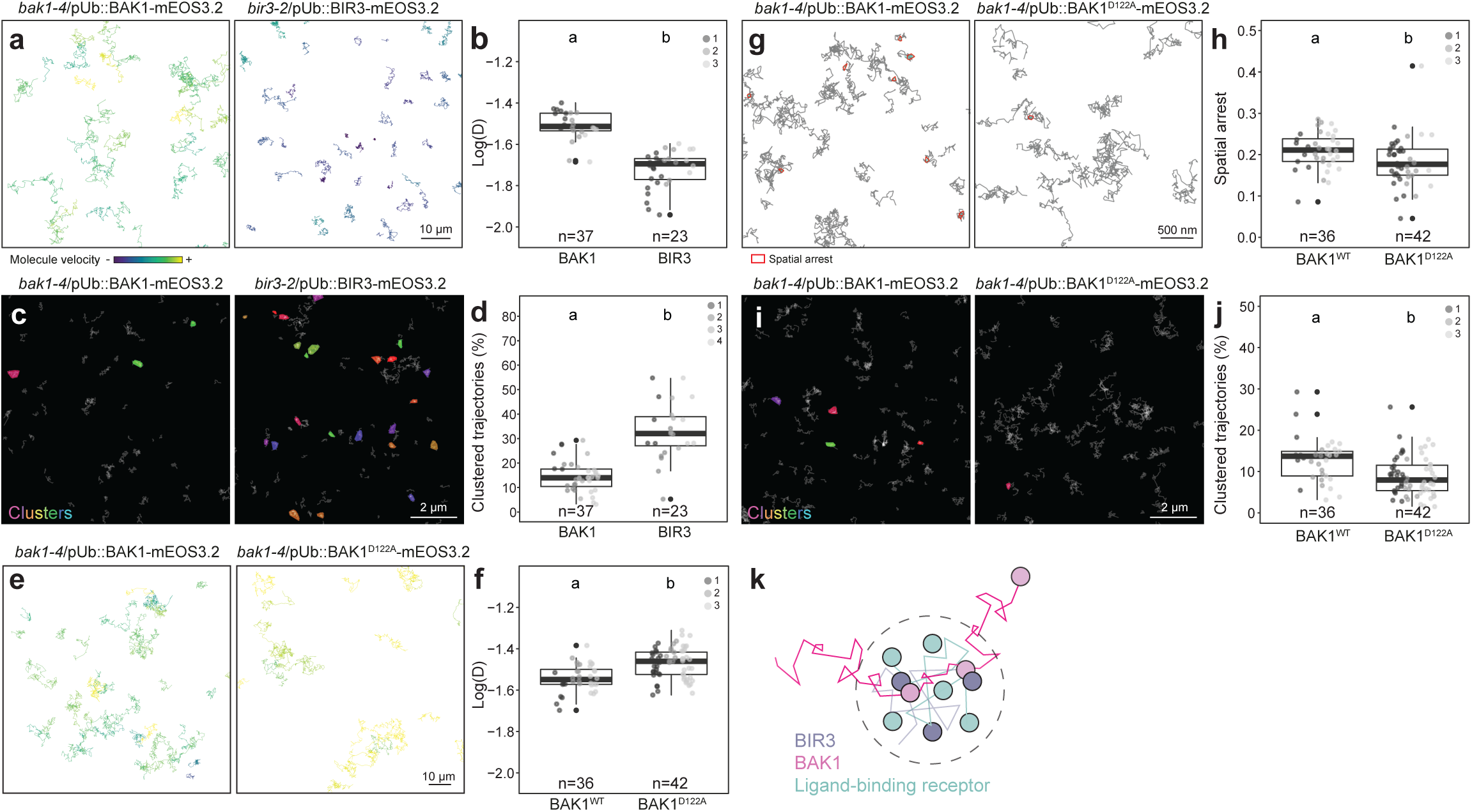
BIRs regulates BAK1 plasma membrane nanoscale dynamics. **a** and **e**, Representative BAK1-mEOS3.2 and BIR3-mESO3.2 (**a**) or BAK1-mEOS3.2 and BAK1^D122A^-mEOS3.2 (**e**) trajectories, long-term single molecule imaging, colors indicate velocity. **b** and **f**, Quantification of instantaneous diffusion coefficient (D). Each data point represents the average log(D) value obtained per cell (n), colors indicate independent experiments. **c** and **i**, Representative images of BAK1-mEOS3.2. and BIR3-mEOS3.2 (**c**) or BAK1-mEOS3.2 and BAK1^D122A^-mEOS3.2 (**i**) trajectories nanoclustering. **d** and **j**, Quantification of clustered trajectories. Each data point represents the percentage of clustered trajectories per cell (n), colors indicate independent experiments. **g**, Representative images of BAK1-mEOS3.2 and BAK1^D122A^-mEOS3.2 spatial arrest. **h**, Quantification of BAK1-mEOS3.2 and BAK1^D122A^-mEOS3.2 spatial arrests. Each data point represents the average number of spatial arrests per trajectory obtained per cell (n), colors indicate independent experiments. Conditions sharing a letter are not significantly different in pairwise Wilcoxon with Bonferroni correction (p>0.05). **k**, Working model: pre-formed ligand-binding RK nanodomains contain accessory RKs such as BIR3.

We next set to reconstitute the plasma membrane nanoscale dynamics of ligand-induced complex formation *in silico* using particle-based simulations (Fig. 4a)^53^. We used the diffusion parameters extracted from single molecule imaging and known ECD-ECD binding kinetics^13,54^ to parametrize our model. These simulations closely recapitulated the nanoscale behavior of single receptor molecules (Extended Data Fig. 8a-c). Interestingly, we observed that introducing BIR3 in these simulations promotes the existence of a dynamic pool of BAK1 in the vicinity of FLS2 nanodomains (Fig. 4b-d) and flg22-induced complex formation (Fig. 4e), suggesting that BIR3 has a positive role in regulating ligand-induced complex formation. In agreement, we observed that loss of *BIR3* (*bir3-2* allele^14^ and *bir3-c* CRISPR mutant (Extended Data Fig. 7)) led to a decrease in flg22-induced FLS2-BAK1 complex formation in Arabidopsis seedlings (Fig. 4g; Extended Data Figure 9a-c).

**Figure 4.**
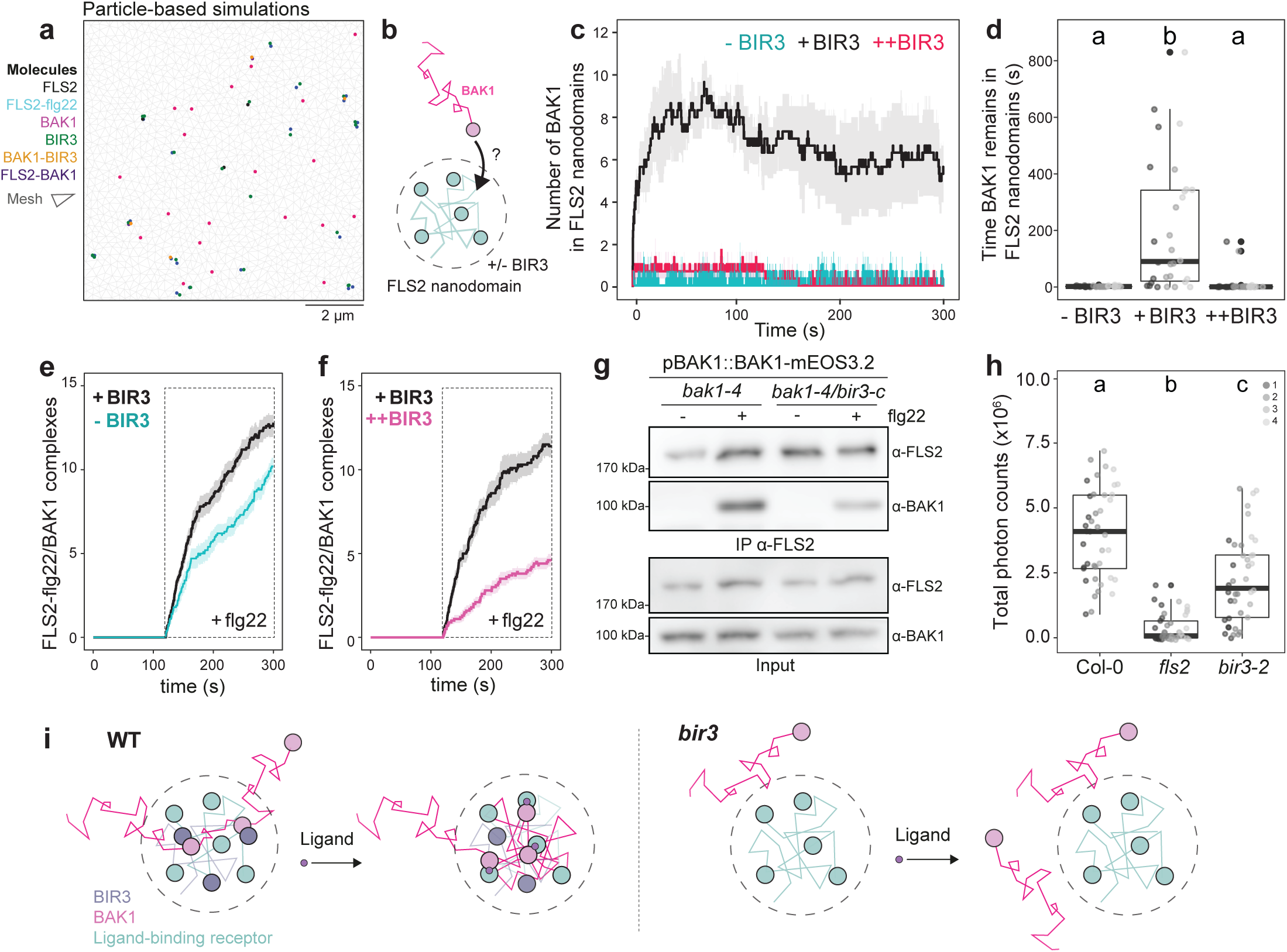
BIR3 maintains a dynamic pool of BAK1 close to FLS2 for ligand-induced complex formation. **a**, Snapshot of a particle-based simulation. **b,** Schematic depicting of the monitoring of BAK1 occurrence in FLS2 nanodomain in presence and absence of BIR3 in particle-based simulations. **c**, Number of BAK1 molecules present in FLS2 nanodomains over time. Data show mean ± se of 3 independent simulations. **d**, Duration of BAK1 occurrence in FLS2 nanodomains. Data points correspond to individual FLS2 nanodomain entry events, colors indicate independent simulations. **e-f,** Analysis of FLS2-flg22/BAK1 complex formation over time in presence of intermediate levels of BIR3 (+BIR3), no BIR3 (-BIR3) (**e**) or high levels of BIR3 (++BIR3) (**f**). The simulations were run 120 s prior to the addition of flg22 in the system. Data represent mean ± se of 10 independent simulations per scenario. **g**, Immunoprecipitation of FLS2 in *bak1-4*/pBAK1::BAK1-mEOS3.2 or *bir3-c*/*bak1-4*/pBAK1::BAK1-mEOS3.2 CRISPR line after treatment with 20 nM flg22 or corresponding mock solution (H_2_O) for 2 min. Membranes were probed with anti-FLS2 or anti-BAK1. **h**, Reactive oxygen species production (ROS) in 4-week-old Arabidopsis leaf discs elicited with 100 nM flg22. Data points indicate total photon counts measured in individual leaf discs (n), colors indicate independent experiments. Conditions sharing a letter are not significantly different in pairwise Wilcoxon test with Bonferroni correction (p<0.05). **i**, Working model: In WT conditions BIR3 attracts and maintains a dynamic pool of BAK1 for efficient ligand-induced complex formation. In absence of accessory receptors, such as BIR3, BAK1 is not maintained in the vicinity of ligand-binding receptors, and the likelihood of ligand-induced complexes is reduced.

Corroborating these observations, loss of *BIR3* led to a decrease in flg22-induced MAPK phosphorylation and flg22-induced ROS production (Fig. 4h; Extended Data Fig. 9d-e). These results contrast with a previous report in Arabidopsis^14^, but align with the effects of loss of BIRs in wheat^55^ and in *N. benthamiana*^56^. In addition, in line with previous observations in Arabidopsis root and dark grown hypocotyls^14^, and in rice^57^, we observed that loss of *BIR3* led to reduced eBL-responsiveness in light-grown Arabidopsis hypocotyls (Extended Data Fig. 9f, g). These results together indicate that BIR3 functions as positive regulator of both flg22- and eBL-triggered signaling. Altogether, these observations suggest that BIR3 attracts BAK1 in the vicinity of ligand-binding receptors, providing a dynamic pool of BAK1 available for ligand-induced complex formation with ligand-binding receptors. We next tested the effect of increasing the amount of BIR3 *in silico*. These simulations predict a decrease in the number of BAK1 molecules in the vicinity of FLS2 (Fig. 4c). Further, in agreement with the effect of BIR3 over-expression observed *in vivo*^14^, increasing the number of BIR3 molecules decreases FLS2-BAK1 complexes formed *in silico* (Fig. 4f). In this system, BIR3 predominantly acted as a negative regulator keeping BAK1 away from FLS2 nanodomains. Our data indicate a dual, concentration-dependent, role for the accessory receptor kinase BIR3.

## Discussion

How cell surface receptors dynamically assemble into signaling-competent complexes within the plasma membrane remains a fundamental question. By combining super-resolution microscopy, long-term single molecule imaging and mathematical modeling, our results suggest a nanoscale organizational principle underlying the formation of Arabidopsis LRR-RK complexes. Ligand-binding receptors are pre-organized into nanodomains, while the common co-receptor BAK1 is diffusive and spatially arrested to ligand-binding receptors. The spatial and temporal orchestration of plant LRR-RK and animal cell-surface receptor complexes thus differ. For instance, compared to FLS2 or BRI1, the human epidermal growth factor receptor (EGFR) and type-I interferon receptors are diffusive and undergo ligand-induced dimerization and spatial stabilization^38,58^. Our experimental and *in silico* analyses converge in describing receptor kinases as an organized and predictable system in which BIR3 maintains a dynamic pool of available BAK1 in the vicinity of ligand-binding receptors for ligand-induced complex formation and signaling (Fig. 4i). We thus propose that accessory receptors, such as BIR3, act as spatial regulators for co-receptor availability. This reconciles previously conflicting genetic observations made in several plant species^14,55–57^ and highlights how subtle changes in accessory receptor abundance can tune ligand responsiveness. Collectively, our findings suggest that ligand-induced complex formation is a deterministic process defined by the relative spatial positioning of individual signaling and regulatory components at the nanoscale rather than purely stochastic diffusion-driven encounters. In this framework, signaling competence emerges from the pre-established spatial architecture of the plasma membrane and may reflect a general organizational principle of signaling systems.

## Acknowledgments

We thank all members of the NanoSignaling Laboratory for fruitful discussions and comments on the manuscript. We thank Niko Geldner for providing the Col-0/pBRI1::BRI1-GFP, Sacco de Vries for providing the Col-0/pBRI1::BRI1-GFP/pBAK1::BAK1-mCherry and Col-0/pBAK1::BAK1-mCherry lines, and Birgit Kemmerling for providing the *bir3-2* mutant. Bio-imaging was performed at the ZMBP microscopy facility. We thank Sven zur Oven-Krockhaus for maintaining the VA-TIRFM. We thank Birgit Kemmerling, Thorsten Nürnberger and Kyle Bender for comments on the manuscript. This research was supported by the Deutsche Forschungsgemeinschaft (DFG) grants (A09-SFB1101 and B01-TRR356) to J.G., the European Molecular Biology Organization (postdoctoral fellowship EMBO LTF 438-2018, to J.G.), the European Research Council under the European Union’s Horizon 2020 research and innovation programme grant agreement no. 773153 (project ‘IMMUNO-PEPTALK’ to C.Z.), the University of Zürich (to C.Z.), the University of Tübingen (to J.G.) and the Technical University of Munich (to J.G.).

## Figures legends

**Extended Data Figure 1.**
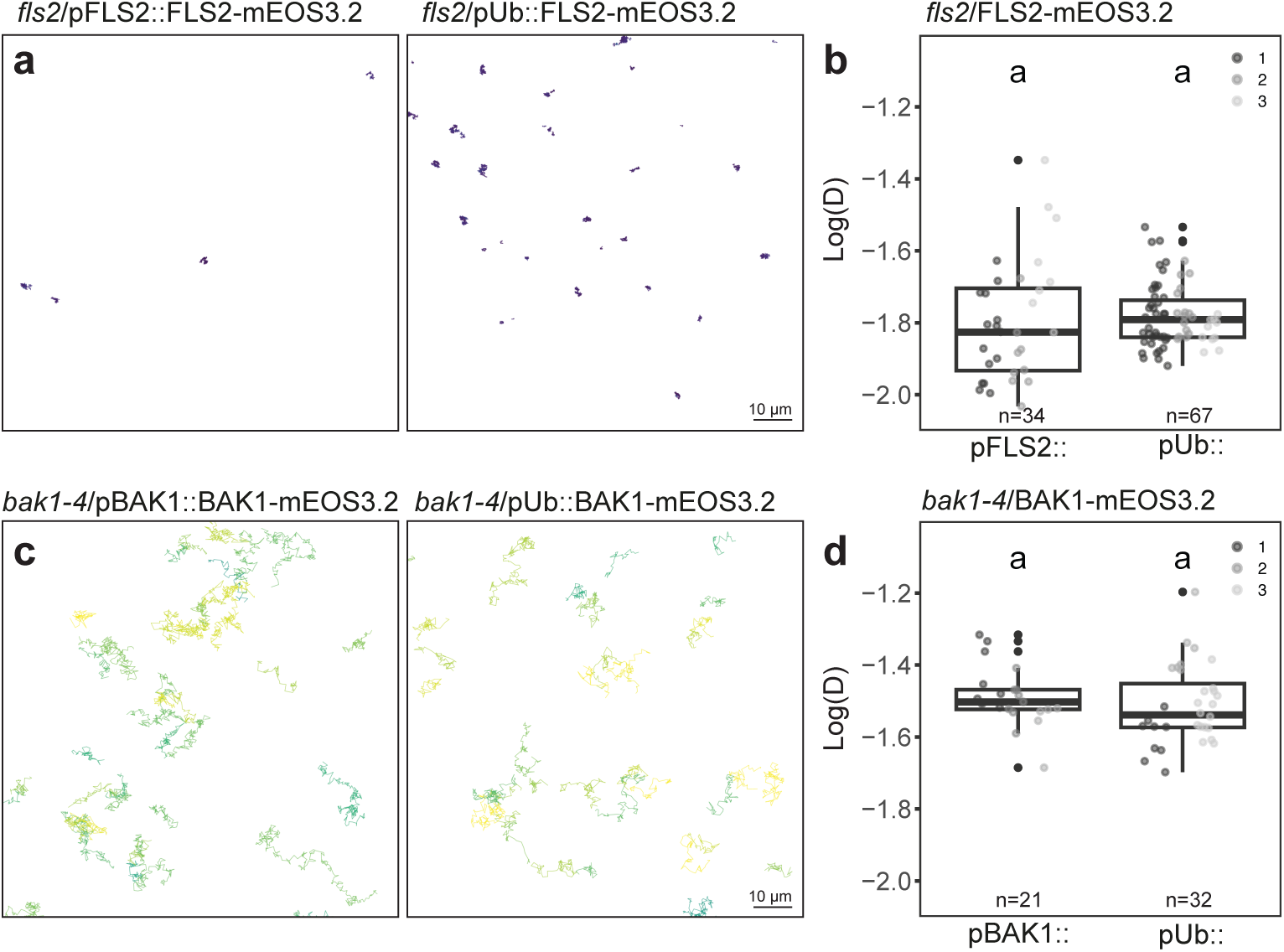
Analysis of FLS2 and BAK1 diffusion in Arabidopsis. **a** and **c**, Representative single molecule trajectories of FLS2-mEOS3-2 and BAK1-mEOS3.2, long-term single molecule imaging in Arabidopsis hypocotyl epidermis, colors indicate velocity. **b** and **d**, Quantification of instantaneous diffusion coefficient (D). Each data point represents the average log(D) value obtained per cell (n). Conditions sharing a letter are not significantly different in pairwise Wilcoxon test with Bonferroni correction (p<0.05).

**Extended Data Figure 2.**
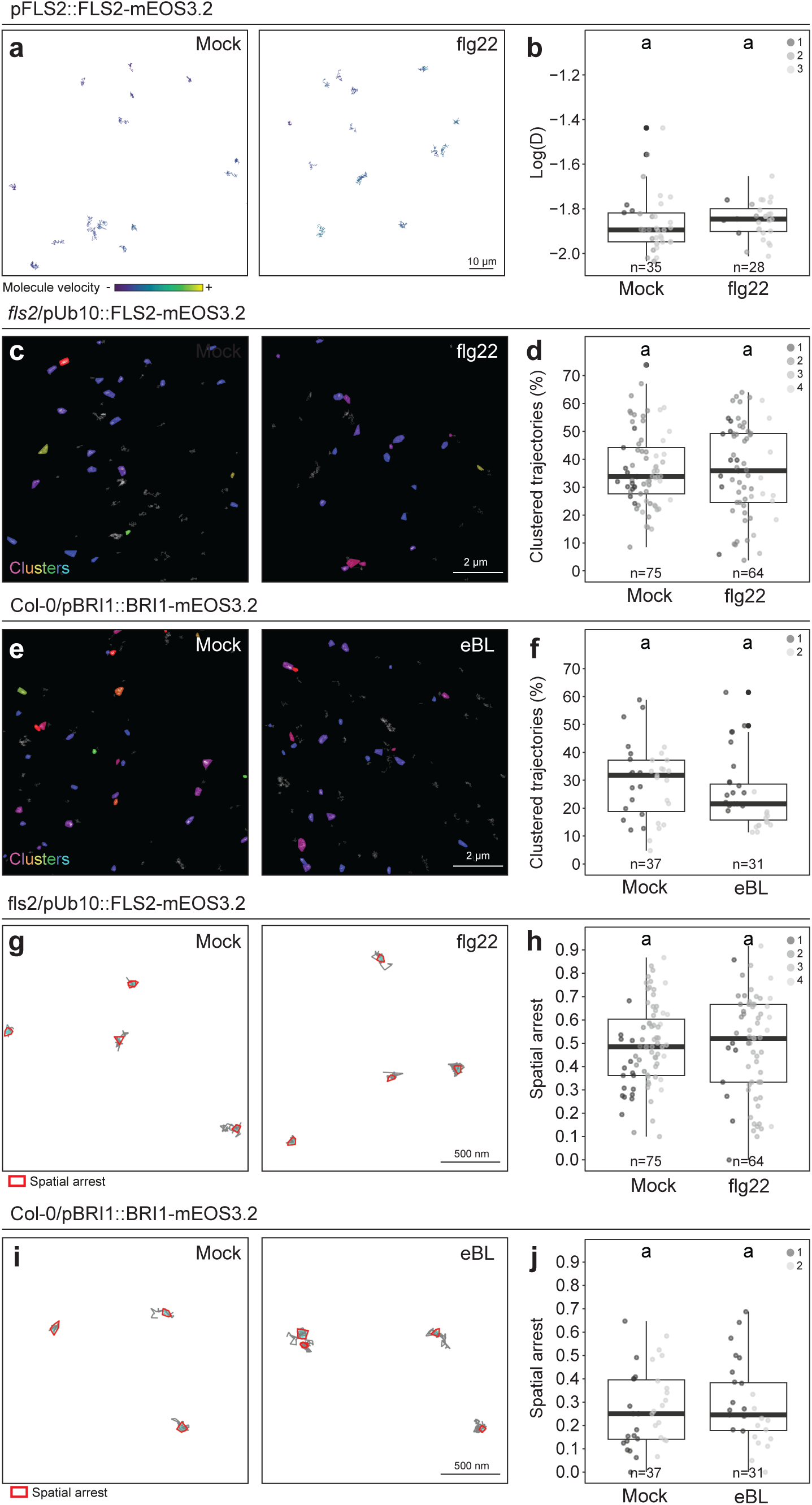
Analysis of FLS2 and BRI1 diffusion and organization in presence and absence of exogenous ligand treatment. **a,** Representative single molecule trajectories of FLS2-mEOS3.2 in presence of 1 μM flg22 or corresponding mock control (H_2_O) in *N. benthamiana*. **b**, Quantification of instantaneous diffusion coefficient (D) within 10 min of treatment. Each data point represents the average log(D) value obtained per cell (n), colors indicate independent experiments. **c, e, g, i,** Representative images of FLS2-mEOS3.2 (**c, g**) or BRI1-mEOS3.2 (**e, i**) single molecule nanoclustering (**c, e**) or trajectories spatial arrest (**g, i**) in Arabidopsis hypocotyl in presence of 1 µM flg22 (**c, g**) or 1 µM eBL (**e, i**) or the corresponding mock controls (H_2_O or EtOH). **d** and **f**, Quantification of FLS2-mEOS3.2 (**d**) or BRI1-mEOS3.2 (**f**) of clustered trajectories within 10 min of treatment. Each data point represents the percentage of clustered trajectories per cell (n), colors indicate independent experiments. Conditions sharing a letter are not significantly different in pairwise Wilcoxon with Bonferroni correction (p>0.05). **h** and **j,** Quantification of FLS2-mEOS3.2 (**h**) or BRI1-mEOS3.2 (**j**) spatial arrest within 10 min of treatment. Each data point represents the average number of spatial arrests per trajectory obtained per cell (n), colors indicate independent experiments

**Extended Data Figure 3.**
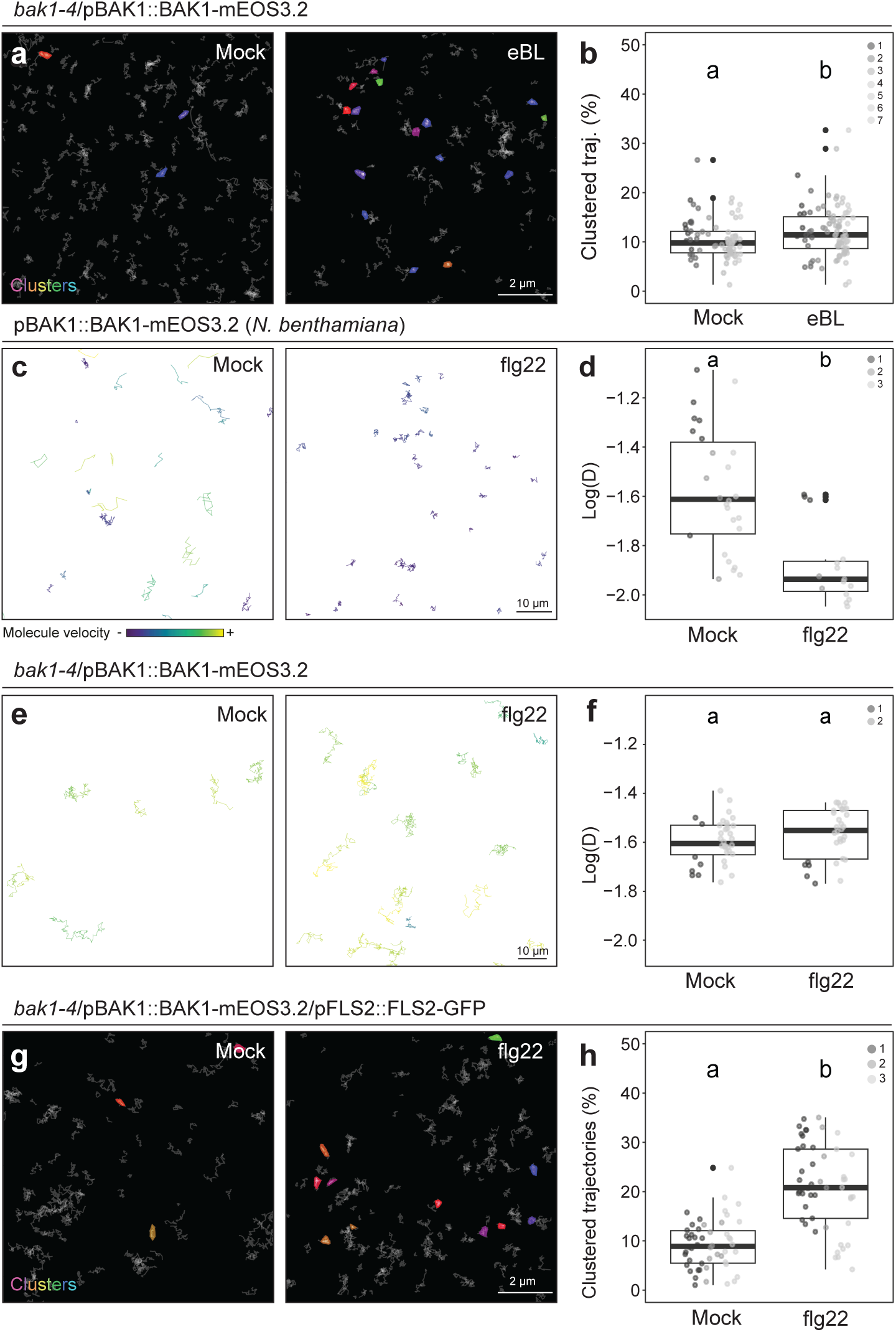
Analysis of BAK1 diffusion and organization in presence and absence of exogenous ligand treatment. **a** and **g**, Representative images of BAK1-mEOS3.2 trajectories nanoclustering in Arabidopsis hypocotyl in presence of 1 µM eBL (**a**) or 1 µM flg22 (**g**) or the corresponding mock controls (EtOH or H_2_O) in *bak1-4* (**a**) and *bak1-4*/pFLS2::FLS2-GFP (**g**). **b** and **h**, Quantification of clustered trajectories within 10 min of treatment. Each data point represents the percentage of clustered trajectories per cell (n), colors indicate independent experiments. **c** and **e**, Representative BAK1-mEOS3.2 trajectories in presence of 1 µM flg22 or the corresponding mock control (H_2_O) expressed in *N. benthamiana* leaves (**c**) or stably expressed in Arabidopsis hypocotyl (**e**). **d** and **f,** Quantification of instantaneous diffusion coefficient (D) within 10 min of treatment. Each data point represents the average log(D) value obtained per cell (n), colors indicate independent experiments. Conditions sharing a letter are not significantly different in pairwise Wilcoxon test with Bonferroni correction (p<0.05).

**Extended Data Figure 4.**
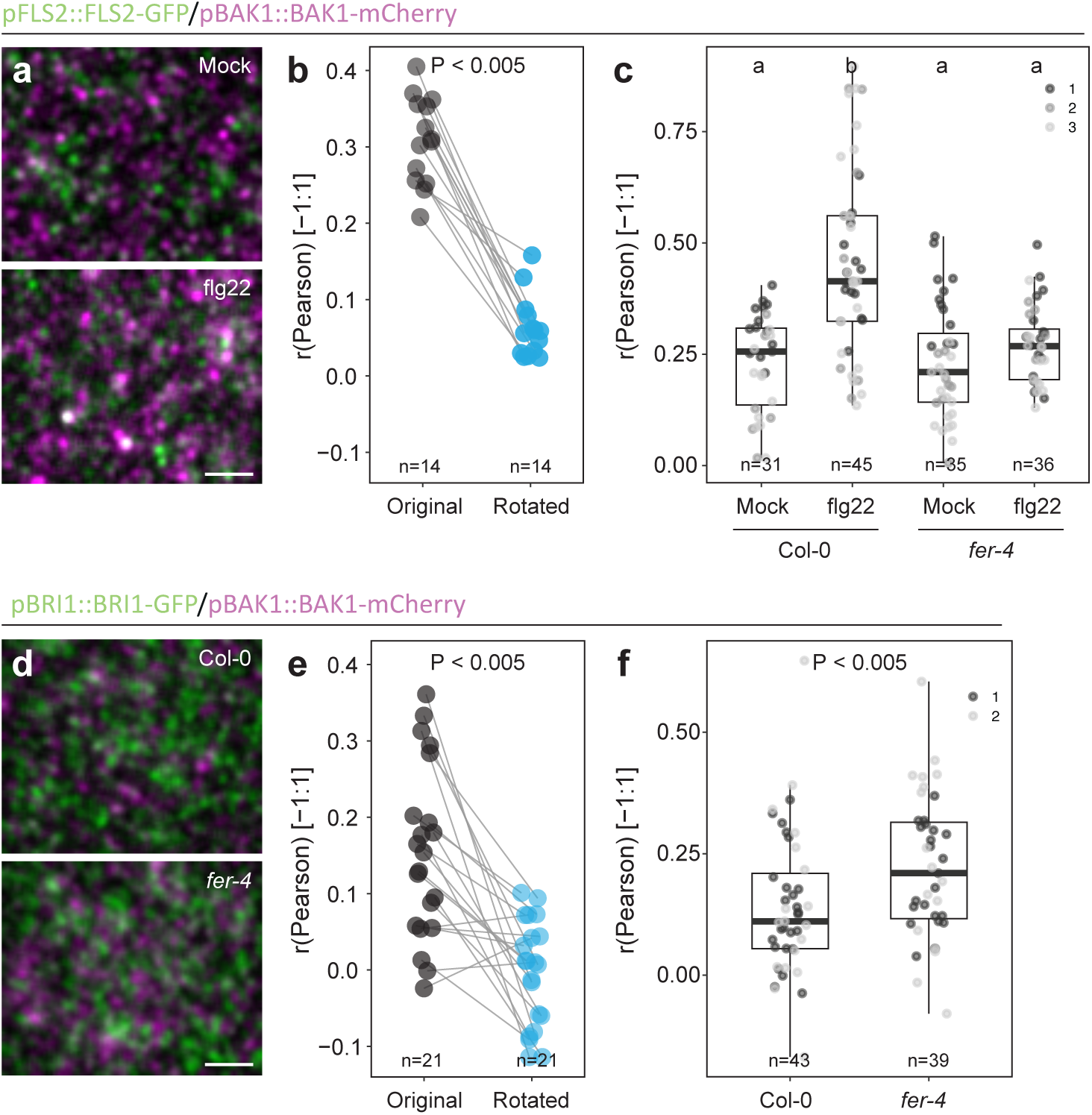
Analysis of FLS2-BAK1 and BRI1-BAK1 co-localization. **a** and **d**, Representative VA-TIRFM images of FLS2-GFP and BAK1-mCherry (**a**), and BRI1-GFP and BAK1-mCherry. **b** and **e**, Quantification of the co-localization between FLS2-GFP and BAK1-mCherry (**b**) and between BRI1-GFP and BAK1-mCherry (**e**). Each data point represents Pearson correlation coefficient for individual cells (n), lines connect raw and corresponding control rotated (90° mCherry) images. **c** and **f**, Quantification of the co-localization between FLS2-GFP and BAK1-mCherry (**c**) and between BRI1-GFP and BAK1-mCherry (**f**), in presence of 1 µM flg22 or the corresponding mock control (H_2_O) (**c**) or control condition (f) in WT Col-0 or *fer-4*. Each data point represents the average Pearson correlation coefficient for individual cells (n), colors indicate independent experiments. Conditions which do not share a letter are significantly different in pairwise Wilcoxon test with Bonferroni p-value adjustment (p< 0,05).

**Extended Data Figure 5.**
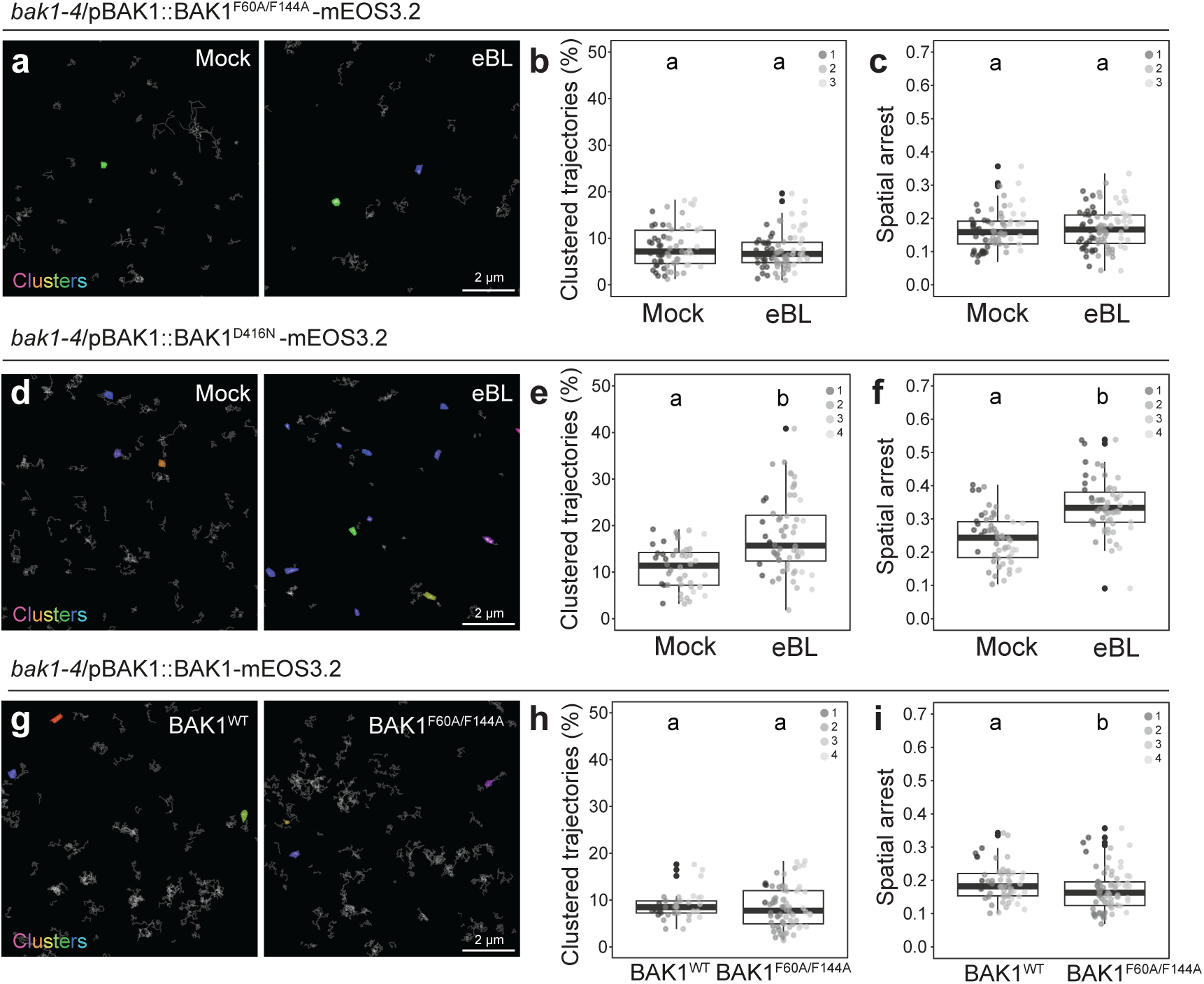
Analysis of the organization of BAK1 mutants in presence and absence of exogenous eBL treatment. **a, d** and **g**, Representative single molecule trajectories clustering of BAK1^F60AF144A^-mEOS3.2 (**a**), BAK1^D416N^-mEOS3.2 (**d**) or BAK1^WT^-mEOS3.2 and BAK1^F60AF144A^-mEOS3.2 (**g**) in presence of 1 µM eBL (**a** and **d**) or the corresponding mock control (EtOH) (**a, d** and **g**). **b, e** and **h**, Quantification clustered trajectories within 10 min of treatment. Each data point represents the percentage of clustered trajectories per cell (n), colors indicate independent experiments. **c, f** and **i**, Quantification of BAK1^F60AF144A^-mEOS3.2 (**c**), BAK1^D416N^-mEOS3.2 (**f**) or BAK1^WT^-mEOS3.2 and BAK1^F60AF144A^-mEOS3.2 (**i**) spatial arrest within 10 min of treatment. Each data point represents the percentage of clustered trajectories per cell (n), colors indicate independent experiments. Conditions sharing a letter are not significantly different in pairwise Wilcoxon test with Bonferroni correction (p<0.05).

**Extended Data Figure 6.**
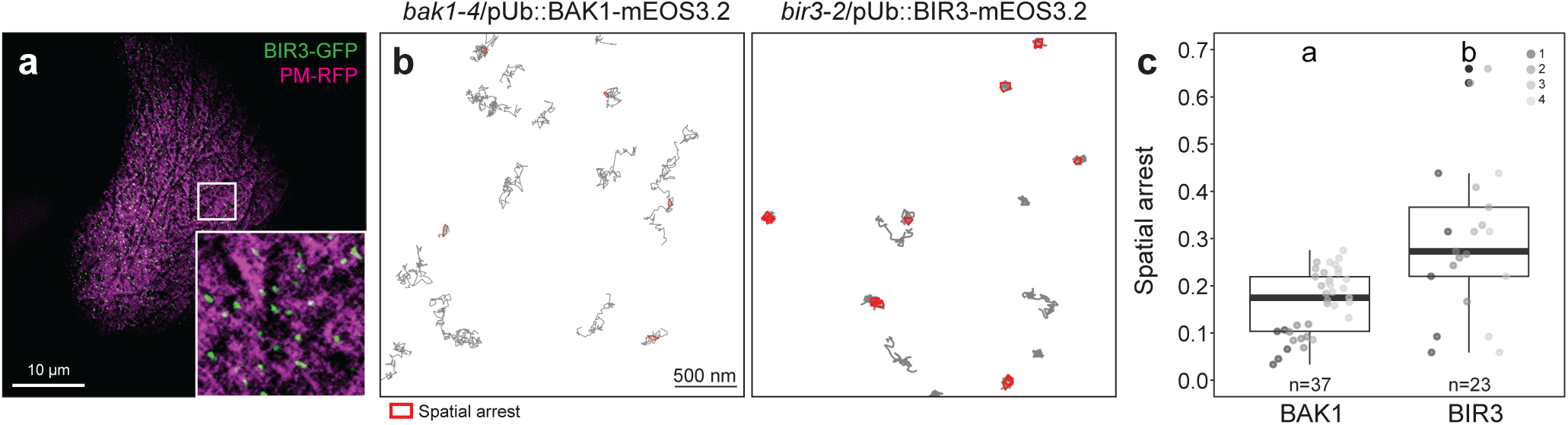
Analysis of BIR3 organization and diffusion. **a**, Representative eSRRF images of BIR3-GFP control plasma membrane (PM) protein expressed in *N. benthamiana*. **b**, Representative single molecule trajectories of BAK1-mEOS3.2 and BIR3-mESO3.2, long term single ^molecule^ imaging in Arabidopsis hypocotyl. **c**, Quantification of spatial arrests. Each data point represents the average number of spatial arrests per trajectory obtained per cell (n), colors indicate independent experiments. Conditions sharing a letter are not significantly different in pairwise Wilcoxon test with Bonferroni correction (p<0.05).

**Extended Data Figure 7.**
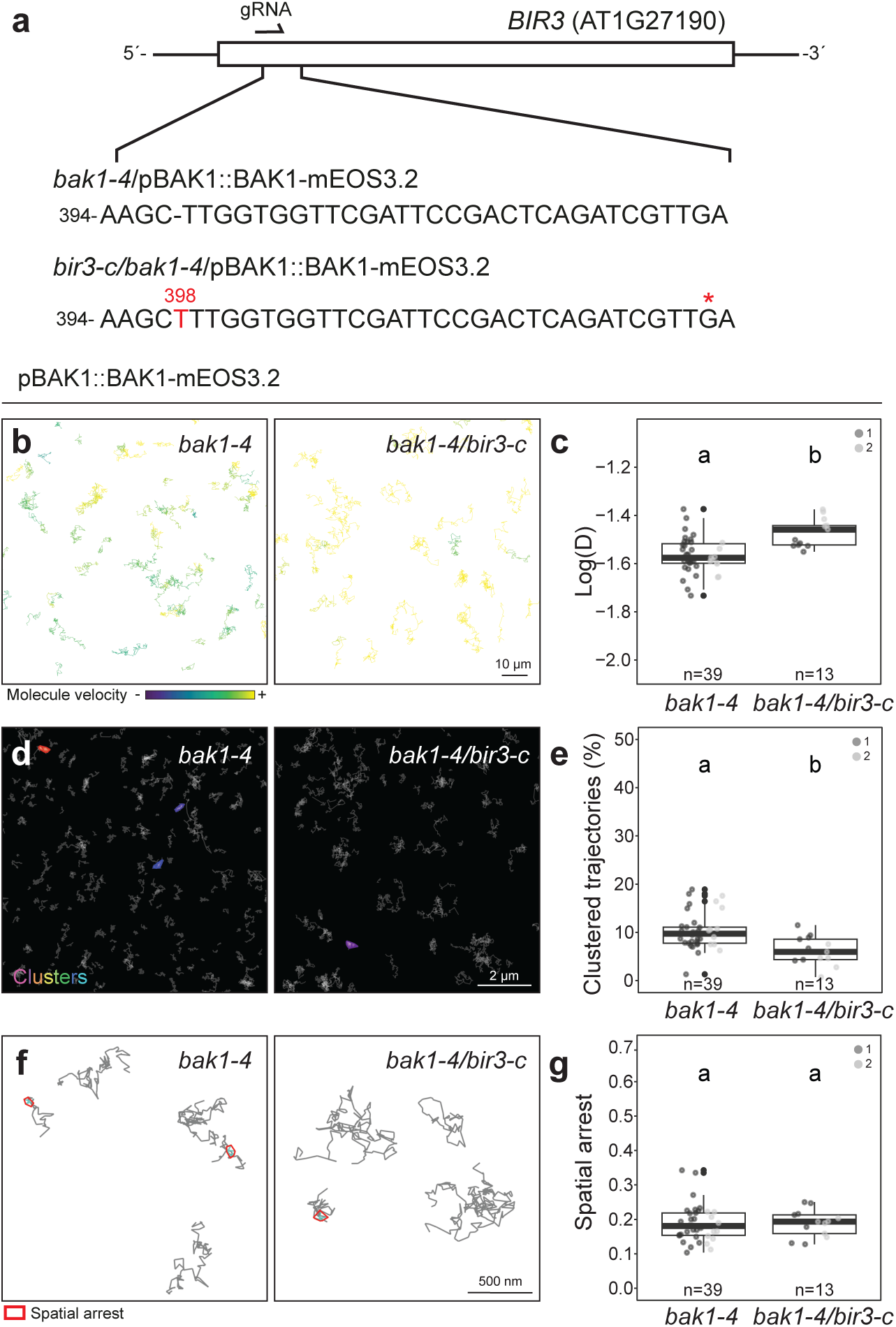
Analysis of BAK1 diffusion and organization in a *BIR3* CRISPR mutant. **a**, Schematic describing the *bir3-c*/*bak1-4*/pBAK1::BAK1-mEOS3.2 CRISPR line. The number of base pairs from the start codon is indicated. Red letters represent insertions observed in gDNA sequencing. Asterisks indicate early stop codons. **b, d, f**. Representative trajectories of BAK1-mEOS3.2, long-term single molecule imaging in *bak1-4* or *bak1-4*/*bir3-c* Arabidopsis hypocotyls. **c**, Quantification of instantaneous diffusion coefficient (D). Each data point represents the average log(D) value obtained per cell (n), colors indicate independent experiments. **e**, Quantification of clustered trajectories. Each data point represents the percentage of clustered trajectories per cell (n), colors indicate independent experiments. **g**, Quantification of spatial arrests. Each data point represents the average number of spatial arrests per trajectory ^obtained^ per cell (n), colors indicate independent experiments. Conditions sharing a letter are not significantly different in pairwise Wilcoxon test with Bonferroni correction (p>0.05).

**Extended Data Figure 8.**
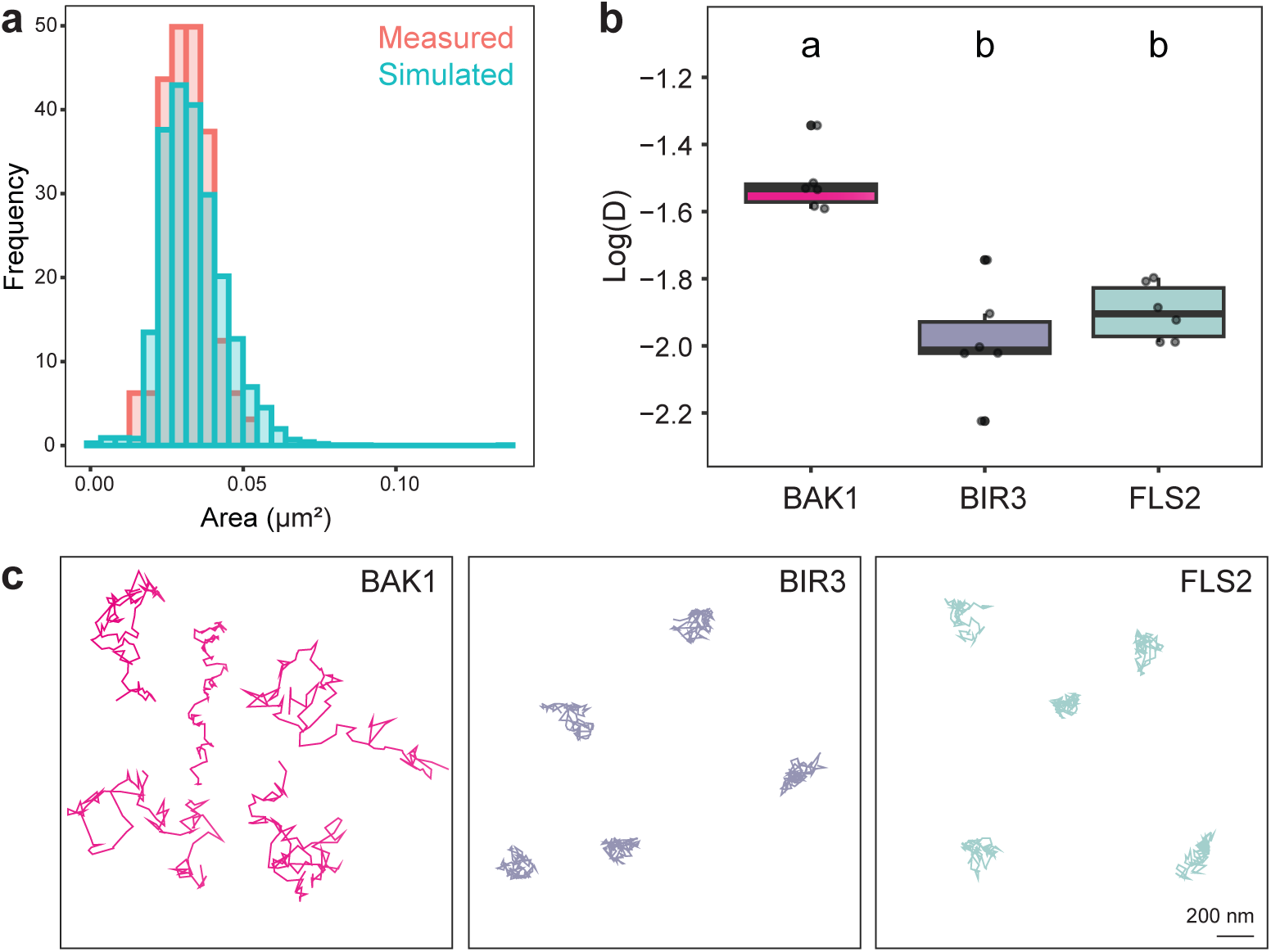
Particle-based simulation recapitulates native protein behavior. **a**, Quantification of the area of simulated FLS2 confinement zones (blue) and measured FLS2-mEOS3.2 nanoclusters observed by spt-PALM and analysed by NASTIC (red). **b**, ^Quantification^ of instantaneous diffusion coefficient (D) of simulated BAK1, FLS2 and BIR3 molecules. Conditions sharing a letter are not significantly different in pairwise Wilcoxon test with Bonferroni correction (p<0.05). **c**, Representative single molecule tracks of simulated BAK1, FLS2 and BIR3 molecules.

**Extended Data Figure 9.**
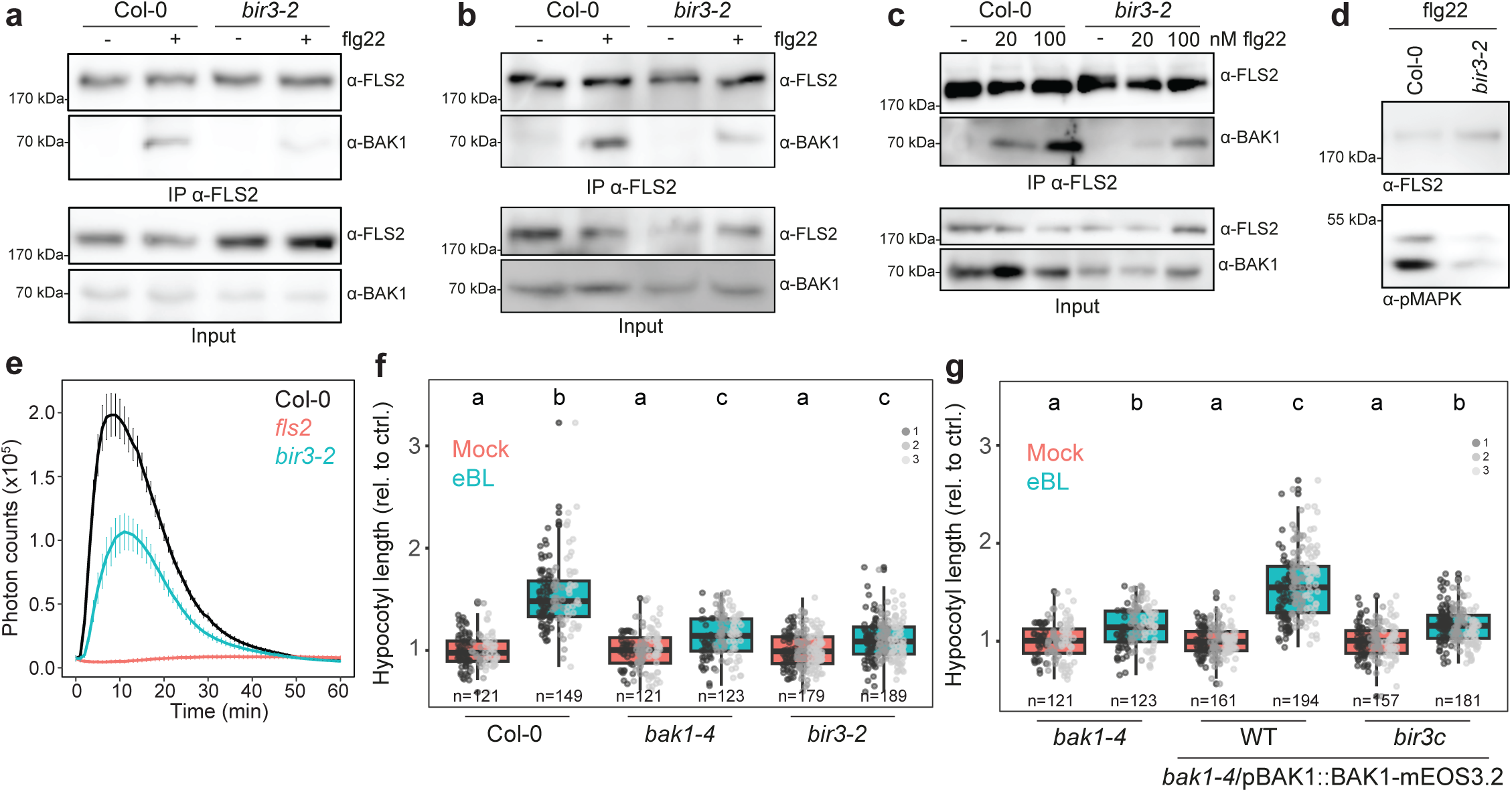
Analyses of eBL and flg22 responsiveness in *BIR3* loss-of-function mutants. **a**, **b**, and **c** Immunoprecipitation of FLS2 in Col-0 or *bir3-2* after treatment with 100 nM flg22 (a, b, c) or corresponding mock solution (H_2_O) for 5 min (**a, b, c**) or 20 nM flg22 and corresponding mock solution (H_2_O) for 2 min (**c**). Membranes were probed with anti-FLS2 or anti-BAK1. **c**d Western blot analysis of MAPK phosphorylation in Arabidopsis seedlings treated for 5 min with 100 nM flg22. Membranes were probed with anti-FLS2 or anti-pERK42/44 (pMAPK) antibodies. The same observations were made in 3 independent experiments. **e**, Time course analysis of ROS production in 4-week-old Arabidopsis leaf discs treated with 100 nM flg22, values are mean ± se of 4 independent experiments. **f** and **g**, Quantification of the hypocotyl length of 5-day-old Arabidopsis seedlings grown on half MS medium containing 200 nM epi-Brassinolide (eBL) or corresponding mock control solution (EtOH). Data points indicate measurements of the length of individual hypocotyl relative to mock for each genotype, colors indicate independent experiments Conditions sharing a letter are not significantly different in pairwise Wilcoxon test with Bonferroni correction (p<0.05).

## Material and methods

### Plant materials and growth

*Arabidopsis thaliana* ecotype Columbia (Col-0) was used as WT control. Col-0/pFLS2::FLS2-GFP^59^, *bak1-4*^11^, Col-0/pBRI1::BRI1-GFP^25^, *fls2c*^60^, *bir3-2*^14^, Col-0/pBRI1::BRI1-GFP/BAK1-mCherry^20^, *bak1-4*/pBAK1::BAK1-GFP^26^, Col-0/pBRI1::BRI1-mEOS3.2^61^, were previously described. Col-0/pFLS2::FLS2-GFP^59^, was crossed with Col-0/pBAK1::BAK1-mCherry^62^ to obtain Col-0/pFLS2::FLS2-GFP/pBAK1::BAK1-mCherry. The following stable Arabopidopis lines *fls2*/pUb::FLS2-mEOS3.2, *fls2*/pFLS2::FLS2-mEOS3.2, *bak1-4/*pBAK1::BAK1-mEOS3.2, *bak1-4/*pUb::BAK1-mEOS3.2, *bak1-4/*pBAK1::BAK1^D416N^-mEOS3.2, *bak1-4/*pBAK1::BAK1^F60A/F144A^-mEOS3.2, *bak1-4/*pUb::BAK1^D122A^-mEOS3.2 and *bir3-2*/pUb::BIR3-mEOS3.2 were generated by the transforming corresponding knock-out backgrounds via floral dipping^63^. The transformants were selected on hygromycin (25 mg/L) to obtain homozygous mono insertional lines and analyzed by western blotting using anti-BAK1 or anti-FLS2 antibodies. Col-0/pFLS2::FLS2-GFP/pBAK1::BAK1-mCherry and Col-0/pBRI1::BRI1-GFP/BAK1-mCherry were crossed with *fer-4*^64^ to obtain *fer-4*/pFLS2::FLS2-GFP/pBAK1::BAK1-mCherry and *fer-4*/pFLS2::FLS2-GFP/pBAK1::BAK1-mCherry. Col-0/pFLS2::FLS2-GFP was crossed with *bak1-4*/pBAK1::BAK1-mEOS3.2 to obtain *bak1-4*/pFLS2::FLS2-GFP/pBAK1::BAK1-mEOS3.2. *bir3-c/bak1-4*/pBAK1::BAK1-mEOS3.2 was obtained by transforming *bak1-4/*pBAK1::BAK1-mEOS3.2 with CRISPR-Cas9 constructs carrying a gRNA targeting *BIR3*. Transformants were selected based on seed coat fluorescence, sequenced and T3 Cas9i-negative homozygous *BIR3* mutants were selected for functional analyses. To analyse flg22-induced ROS production, plants were grown in individual pots at 21 °C with a 10-h light photoperiod in environmentally controlled growth rooms. For seedlings-based assays, seeds were surface-sterilized by incubating them in 0.1 % Tween20 in 70 % EtOH for 10 min, following 70 % EtOH for 10 min and 100% EtOH for 1 min. Seeds were stratified for 2 d in the dark at 4 °C and grown on half Murashige and Skoog (MS) media supplemented with vitamins, 1 % sucrose and 0.8 % agar at 22 °C and a 16-h light photoperiod.

### Molecular cloning, plant transformation and CRISPR-Cas9-mediated mutagenesis

FLS2, BAK1 and BIR3 coding sequences and FLS2 and BAK1 promotors were obtained as synthetic fragments (Genewiz) and cloned into the level 1 Golden Gate plasmid pUC57_BB03 using Bpil^65,66^. The plasmid encoding for mEOS3.2 was previously described^32^. Nucleic acid fragments were assembled in the level 2 Golden Gate plasmid (BB24_LIIβ F 3-4)^66^. *BIR3* gRNA for CRISPR Cas-mediated editing was designed using CHOPCHOP (https://chopchop.cbu.uib.no/) and inserted into pAGM55261 using Golden Gate cloning^65^. All plasmids were verified by Sanger sequencing. The binary plasmids were transferred into the *Agrobacterium tumefaciens* strain GV3101 for flower dip transformation of Arabidopsis or transient transformation of *Nicotiana benthamiana* leaves.

### Brassinosteroid-sensitivity assay

Brassinosteroid sensitivity assay was performed as previously described in^43^. Briefly, seeds were surface-sterilized and individually placed in line on square Petri dishes containing half MS 1 % sucrose, 0.8 % phytoagar, supplemented with 200 nM epi-brassinolide (Sigma, Cat#E1641) or corresponding control solution (EtOH). The seeds were stratified at 4 °C for 2 d and then placed vertically in a growth chamber for 5 d. Hypocotyl length were measured using Fiji^67^.

### Co-immunoprecipitation

Sample preparation and immunoprecipitation were performed as previously described^68^. Briefly, seedlings were grown on half MS media for 5 d and transferred to 6-well plates containing liquid half MS to be cultivated for ten more days. Subsequently, seedlings from 10 wells were gathered and let rest for 1 h in liquid half MS. Seedlings were treated for 4 min with either 20 nM or 100 nM flg22 or mock (distilled sterile water) and flash frozen in liquid nitrogen. For protein extraction, tissues were grinded in liquid nitrogen and proteins were isolated in 50 mM Tris-HCl pH 7.5, 150 mM NaCl, 10 % glycerol, 5 mM dithiothreitol, 1 % protease inhibitor cocktail (Sigma-Aldrich), 2 mM Na2MoO4, 2.5 mM NaF, 1.5 mM activated Na3VO4, 1 mM phenylmethanesulfonyl fluoride, and 0.5 % IGEPAL. For immunoprecipitations, trueblot agarose beads (Rockland, 18-8816-33) were conjugated with FLS2 antibodies (Agrisera, AS12 1857) and incubated with the crude protein extracts for 3 to 4 h at 4 °C. Subsequently, beads were washed three times with a washing buffer (50 mM Tris-HCl pH 7.5, 150 mM NaCl, 1 mM phenylmethanesulfonyl fluoride, 0,1 % IGEPAL) before adding 60 µL of a protein loading solution (6xSDS 10 % DTT) and incubating for 10 min at 90 °C. Analysis was carried out by SDS-PAGE and immunoblotting.

### Immunoblotting

Protein samples were separated in 10 % bisacrylamide gels at 150 V for approximately 2 hours and transferred into activated PVDF membranes at 100 V for 90 min. Immunoblotting was performed with antibodies diluted in blocking solution (5 % fat-free milk in TBS with 0.1 % [v/v] Tween-20). Antibodies used in this study were α-BAK1 (1:5000, Agrisera AS12 1858), α-FLS2 (1:5000, Agrisera AS12 1857) or α-pMAPK (1:5000, pERK42/44 cell signal 9101S). Blots were developed with Pierce ECL/ECL Femto Western Blotting Substrate (Thermo Scientific). The following secondary antibody was used: anti-rabbit IgG (whole molecule)–HRP (A0545, Sigma, dilution 1:10,000).

### Variable angle total internal reflection microscopy (VA-TIRFM)

VA-TIRM imaging was performed as previously described in^32^ Briefly, VA-TIRFM was operated on a custom-build TIRFM set-up^61^ equipped with an 100x objective NA 1.49 (Zeiss, 421190-9800-000), four laser lines (405, 488, 561 and 642 nm), a polychromatic modulator (AOTF, AA OPTO-ELECTRONIC) and a sCMOS camera (Hamamatsu Photonics, ORCA-Flash4.0 V2). To ensure that the target laser power value always corresponds to the irradiance in the sample plane, the power of each laser line was calibrated before each experiment using a PM100D detector (Thorlabs). Five-day-old Arabidopsis seedlings or 3-week-old *N. benthamiana* leave samples (28 to 34 h post Agrobacterium infiltration) were delicately mounted between two coverslips (Epredia 24×50 mm #1) in liquid ½ MS medium and placed on the specimen stage without additional weight. Images were acquired at 20 Hz frame rate (50 ms), 6.66 Hz (150 ms) or 5 Hz (200 ms) for spt-PALM and photochromic reversion imaging conditions.

### Single molecule imaging and analyses

Photochromic reversion and spt-PALM imaging were performed as previously described^32^. Briefly, for photochromic reversion, mEOS3.2 was photoconverted, excited and kept in a prolonged fluorescent state using 1000 µW 561nm laser power, 2-15 µW 405 nm laser power and 1-30 µW 488 nm laser power. The emitted light was collected using 568 LP Edge Basic Long-pass Filter, 584/40 ET Band-pass filters (AHF analysentechnik AG) and recorded from a 51.2 x 51.2 µm region of interest, 100 nm pixel size. Track reconstruction and analysis was carried out as previously described^32^. To ensure bona-fide single-molecule tracking we analyzed frames with relatively low molecule density (ca. 0.1 – 1 molecule per µm^2^). We used the plugin TrackMate^69^ in Fiji^67^ to reconstruct single molecule trajectories. Single particles were segmented frame-by-frame by applying a Laplacian of Gaussian (LoG) filter and estimated particle size of 0.3 μm with a Quality threshold of 30. Individual single particles were localized with sub-pixel resolution using a built-in quadratic fitting scheme. Single-particle trajectories were reconstructed using a simple linear assignment problem^70^ with a maximal linking distance of 0.4 μm and a 2 frames-gap-closing maximum. The coordinates of the single particle trajectories were then further analyzed using a custom-made python script to calculate the mean square displacement (MSD) and diffusion coefficient (D) in batch mode. Only tracks with a minimum of five localizations were kept for analysis. The MSD and diffusion coefficient was calculated based on the first four time points of each trajectory as previously described^71^. Single molecule trajectories clustering was performed using nastic^35^ with the following threshold, time = 20 s, radius factor = 1.2, minimum trajectories = 3, size screen = 0.15 μm. Spatial arrests were analyzed using CASTA, with the default minimum track length of 25 timepoints^32^ (https://pypi.org/project/casta/).

### Enhanced super-resolution radial fluctuations (eSRRF)

Stable transgenic lines expressing BRI1-GFP, BAK1-GFP or FLS2-GFP were imaged by VA-TIRFM as described above at 5 Hz (200 ms) frame rate. Super resolved images were reconstructed using eSRRF^29^ in Fiji^67^. Stream VA-TIRFM acquisitions were processed using the following parameters that were defined by iterative parameter sweep^29^: magnification = 6, radius = 3, sensitivity = 3 and frames = 100. The average (AVG) reconstructions of BRI1-GFP, BAK1-GFP and FLS2-GFP plasma membrane organization were segmented and analysed using Nanonet, a custom-made automated analysis pipeline (https://github.com/NanoSignalingLab/NanoNet), providing nanodomain density and image wide spatial clustering index (iSCI) based on predicted localization density.

### Measurement of reactive oxygen species production

Arabidopsis plants were grown in individual pots at 21 °C with a 10-h light photoperiod in environmentally controlled growth rooms. Leaf discs (4 mm in diameter) were excised from 3- to 4-week-old plants and incubated in 96-well plates containing sterile distilled water overnight. The following day, water was replaced with a solution containing 0.5 µM L-012 and 20 μg/mL horseradish peroxidase (HRP, Sigma Aldrich) and 100 nM flg22 or control (water). Luminescence was measured using a microplate reader (Promega Glomax navigator) over a period of 60 min.

### Particle based simulations

Computational simulations of single particles were performed using the Smoldyn software (version 2.75)^53^, and a custom-made python script. We simulated a 2D space of 10 µm^2^ with periodic boundaries conditions to avoid edge effects. We imposed a triangular mesh with an average compartment size corresponding to the size FLS2 nanoclusters obtained by NASTIC. To simulate FLS2, BAK1 and BIR3 molecular diffusion we provided as input data the instantaneous diffusion coefficient (D) measured by photochromic reversion. For each step and each molecule, a new displacement is randomly chosen from normally distributed possible displacements. The probability of molecule diffusion across compartment boundaries was defined as 0.00001 for FLS2, 0.00001 for BIR3 and 1 for BAK1. To simulate molecule binding kinetics, we used the Kon and Koff of flg22-bound FLS2 to BAK1^54^ and the Kd of BAK1 to BIR3^13^ converted into binding and unbinding radii. Simulations were run using a delta t of 0.0001s to ensure that the spatial resolution was smaller than the geometric features and the characteristic time scale of the system for each possible reaction. The initial location of each molecule was randomized for each simulation. To analyse the effect of BIR3 on BAK1 localization we monitored the number of BAK1 molecules within FLS2 nanodomains every 5 ms for 300 s. To simulate flg22 perception, FLS2 molecules were converted to flg22-bound FLS2 molecules after 120 s and FLS2-flg22/BAK1 complexes were measured every 50 ms for 300 s.

